# Computational modeling of neurovascular coupling at the gliovascular interface

**DOI:** 10.64898/2026.05.15.725343

**Authors:** Florian Dupeuble, Hugues Berry, Audrey Denizot

**Affiliations:** AIstroSight, Inria, Hospices Civils de Lyon, Universite Claude Bernard Lyon 1, F-69603 Villeurbanne, France

## Abstract

A growing number of studies indicate the possible involvement of astrocytes in triggering or modulating neurovascular coupling (NVC), i.e. the local dilation of blood vessels in the brain in response to neuronal activity. Astrocytes possess specialized subcellular compartments, named endfeet, that surround arterioles and capillaries, ideally positioned to mediate NVC. Various vasodilators have been shown to contribute to NVC, such as epoxyeicosatrienoic acid (EET), nitric oxide (NO), or prostaglandin E2 (PGE2), but the precise mechanisms underlying NVC and their variability remain to be fully elucidated. In particular, the involvement of astrocytes in this process is controversial. Recent translatome and proteomics data reveal that astrocytes and in particular endfeet are enriched in the proteins of the PGE2 pathway. However, how the latter could contribute to NVC remains to be characterized. Here, we develop a computational model of astrocyte-mediated NVC that recapitulates these findings and describes Ca^2+^ and PGE2 signaling in astrocytes, NO release by neurons, and arteriole diameter dynamics using ordinary differential equations. The model successfully reproduces the dynamics of arteriole diameter change during hyperemia from *in vivo* neocortical recordings in awake mice. Our simulations suggest that the astrocyte PGE2 pathway could be responsible for the late response of NVC at the arteriolar level. We further observe that PIP2-derived diacylglycerol plays a major role in driving arteriole diameter dynamics in our model, while phosphatidic acid-derived diacylglycerol, which is calcium-dependent, mainly acts as an amplifier of this response. Finally, a spatial implementation of the model using a simplified astrocyte geometry suggests that NVC is more efficient when synaptic stimulation occurs at the endfoot level rather than at other astrocytic compartments. Overall, this computational study suggests a partial role for astrocyte-mediated PGE2 release in NVC and points to astrocyte perivascular processes as sub-compartments that are ideally positioned and equipped to mediate NVC.

**Author summary:** In the brain, the local blood flow is regulated to meet neuronal energy demand by modulating the dilation of neighboring blood vessels. The mechanisms driving this process, known as neurovascular coupling (NVC), remain debated and are likely to differ depending on the physiological context. Recent evidence points to astrocytes, a cell type possessing specialized protrusions called “endfeet”, that envelop the entire brain vascular tree. Contacts between synapses and endfeet have recently been reported, positioning the latter as ideal mediators of NVC. Here, we developed a computational model that simulates the signaling between neurons, astrocytes, and blood vessels. Our model successfully reproduces experimental recordings of blood vessels dilation in the brains of awake mice. Our simulations suggest that a specific signaling pathway in astrocytes, involving a molecule called prostaglandin E2, is a key driver of the late phase of NVC, occurring a few seconds after neuronal activity. Furthermore, our model indicates that the location of the stimulated synapses matters: signals sent to the astrocyte endfeet are particularly effective at controlling blood flow. This work helps clarify the active role of astrocytes in brain blood flow regulation, a process critical for healthy brain function.

## Introduction

Neurovascular coupling (NVC), the local dilation of blood vessels (also called hyperemia) in response to neuronal activity, is essential for brain function. It sustains neuronal activity by meeting the metabolic needs of neurons, and may have broader roles [1]. This coupling underlies the blood-oxygen-level-dependent (BOLD) signal, measured in functional magnetic resonance imaging (fMRI) [2]. Importantly, vascular dysfunctions or neurovascular uncoupling are observed in a wide range of neurological disorders such as Alzheimer’s disease, traumatic brain injury, and multiple sclerosis [3–5]. The mechanisms responsible for these NVC deficiencies are not well understood but could pave the way for novel therapeutic strategies. Glutamate release by active glutamatergic neurons is commonly accepted as the trigger of NVC. Numerous vasodilators downstream glutamate have been suggested to contribute to NVC, such as nitric oxide (NO), prostaglandin E2 (PGE2), or potassium ions, to name a few [4]. A meta-analysis on the various signaling pathways mediating NVC suggested that NO is essential to trigger NVC but identified numerous additional contributors to hyperemia, highlighting the complexity of NVC [6]. NVC dynamics may also vary depending on the type of dilated vessels (arterioles vs capillaries) [7]. Moreover, *in vivo* recordings suggest that NVC dynamics are more complex than simple rise and fall kinetics, and may feature two successive phases, referred to as an “early-phase” and a “late-phase” [8, 9].

Recent studies suggest that neurons are not the only cell types involved in NVC [10]. In particular, astrocytes, glial cells known to interact with neurons and blood vessels, are perfectly positioned to mediate NVC. They possess specialized subcellular compartments, referred to as endfeet, that wrap and cover almost the entire brain vasculature [11]. The involvement of astrocytes in NVC is usually evaluated using their intracellular Ca^2+^ signals as a proxy but remains controversial [12]. Some studies suggest that astrocytes Ca^2+^ elevations are not mandatory to dilate blood vessels [13–15], while others have detected systematic astrocyte Ca^2+^ signals preceding hyperemia, notably in endfeet [16–18]. This apparent discrepancy might result from differences in temporal resolution or from local variability of the neuro-gliovascular unit [19, 20].

Computational models are especially useful in this type of situation, thanks to their ability to disentangle the contributions of the various actors of the system. While most computational models of NVC do not take astrocytes into account [21], several proposals have incorporated them via their Ca^2+^ dynamics [22–27]. However, to the best of our knowledge, the existence of two temporal phases of NVC, early and late, is often ignored in these models or remains phenomenologically described [28], which limits mechanistic interpretability of the underlying biological processes. Moreover, the PGE2 pathway is rarely described in details and is often modeled as being entirely driven by Ca^2+^ signals, without explicit intermediate reactions. Lastly, the astrocyte is most often represented as a single compartment, which fails to account for the extreme subcellular organization of these cells (endfeet, branchlets, leaflets), thereby neglecting their spatial and functional heterogeneity.

To deepen our understanding of the role of astrocytes in NVC, we developed a mean-field computational model of astrocyte-dependent NVC, implemented using Ordinary Differential Equations (ODEs). The model includes an astrocyte and a blood vessel compartment, as well as a phenomenological neuronal nitric oxide (NO) pathway, that accounts for the early phase observed in *in vivo* arteriole diameter recordings [8]. To model the astrocyte compartment, we first analyzed recent translatome and proteomics datasets [29–32], which revealed that all of the enzymes and receptors of the PGE2 pathway are expressed by astrocytes, in particular in endfeet. Our model successfully reproduces the early and late phases of NVC reported *in vivo* [8]. Using extensive simulation campaigns, we show that the model predicts that the astrocyte PGE2 pathway is sufficient to sustain the late phase of NVC but cannot trigger the early phase. Moreover, our results suggest that PGE2-induced arteriole dilation can occur in the absence of astrocyte Ca^2+^ signals, although the latter amplify NVC. Finally, a spatial implementation of the model in a simplified multi-compartmental astrocyte geometry reveals that the influence of the astrocyte PGE2 pathway on NVC strongly depends on the location of the neuronal input relative to the astrocyte tree. In particular, arteriole dilation is predicted to be strongly reduced when neuronal stimulation occurs elsewhere than at the endfoot.

## Materials and Methods

Neurovascular coupling is modeled using ordinary differential equations that couple neuronal activity, astrocyte PGE2 signaling, and vascular diameter dynamics. The complete kinetics scheme of the model is shown in Figure 1. The simulated fluxes are illustrated in supplementary figure S3.

**Fig 1.**
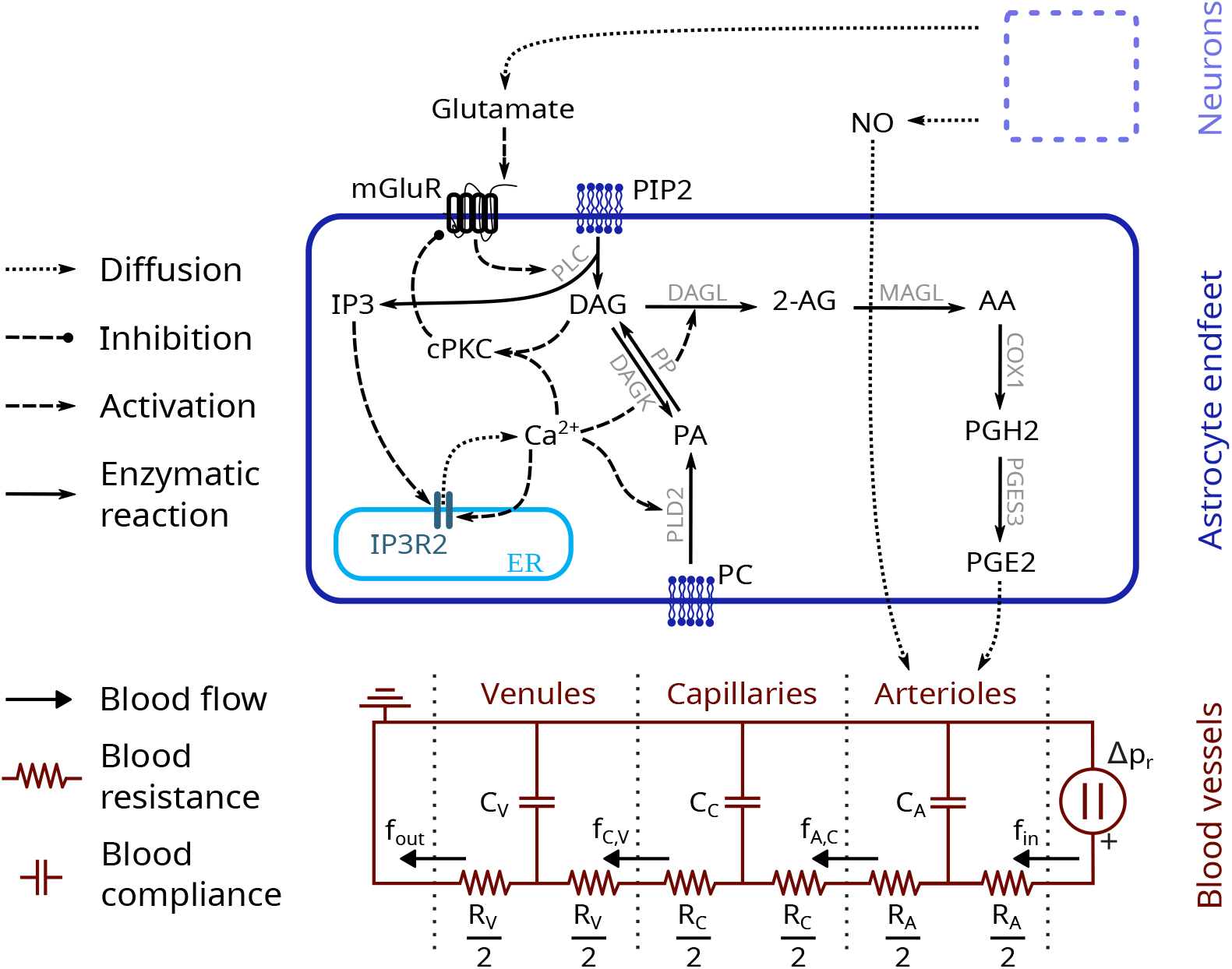
Reaction scheme of the neurovascular coupling model. Neurons release NO and glutamate in the extracellular space. The latter binds to astrocyte mGluRs, triggering calcium-induced calcium release (extended from [33]) and PGE2 production. PGE2, in addition to NO, diffuses to the arterioles, resulting in their dilation. Blood vessels are modeled using an extended 3-compartments Windkessel model [28, 34, 35].

### Neuronal activity model

Neuronal NO release is modeled as a phenomenological component capturing the early and rapid vasodilatory contribution observed experimentally [8] without additional mechanistic complexity. More specifically, it corresponds to a rectangular function, active for the first four seconds of stimulation (see figure 2B).

**Fig 2.**
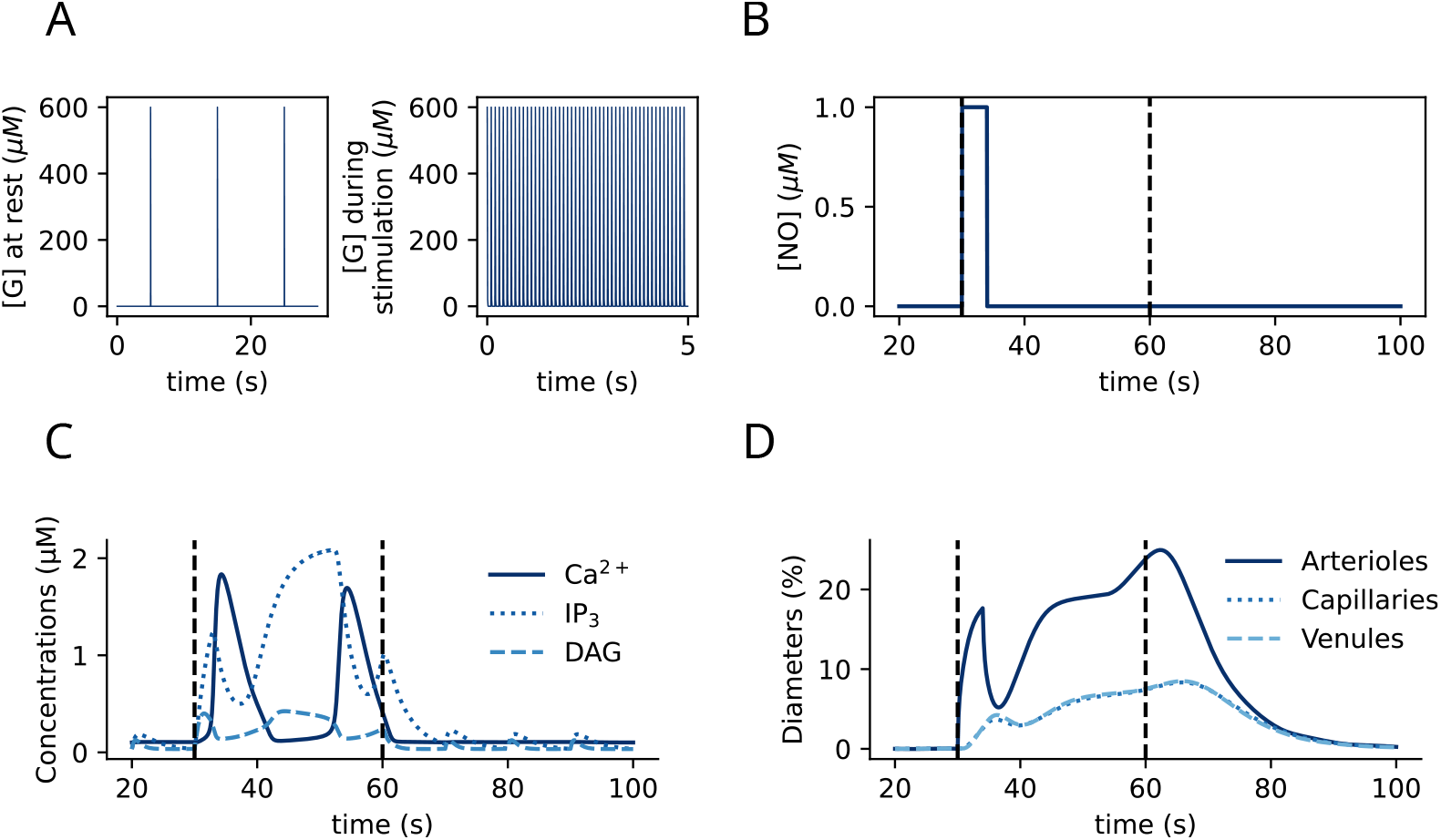
NVC model dynamics. (A) Dynamics of Glutamate concentration [*G*], under resting (left) and stimulation (right) conditions. (B) Dynamics of NO concentration [*NO*], modeled as a rectangular function of time. (C) Representative temporal traces of astrocyte calcium-induced calcium release-related variables : Ca^2+^, IP_3_, and DAG concentrations. (D) Representative temporal traces of diameter changes of arterioles, capillaries, and venules, expressed as a percentage of the steady state diameter. In all panels, black vertical dashed lines delineate the stimulation period.

In line with experimental measurements of glutamate concentration dynamics in the synaptic cleft [36], neuronal glutamate release is modeled as an exponential decay with a decay constant *τ*_G_ = 0.003 s^−1^, combined with discrete presynaptic releases occurring at regularly-spaced spike times *t*_*spike*_:

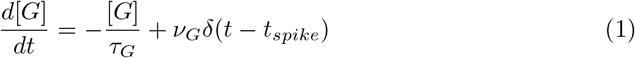

where *G* is the extracellular glutamate concentration, *ν*_*G*_ the glutamate concentration increase occurring at each *t*_*spike*_ and *δ*(*t*) is the Dirac delta.

Spike frequency is set at 0.1 Hz and 10 Hz for rest and during neuronal stimulation, respectively [37]. Figure 2A displays representative temporal traces of [G] when the neurons are at rest or during stimulation.

### Astrocyte endfoot model

#### Identification of the astrocyte reaction scheme

Analysis of recent translatome [29] and proteomics data [30–32] revealed that astrocytes and in particular endfeet are enriched in all proteins of the prostaglandin *E*_2_ (PGE2) pathway except Cyclooxygenase 2 (COX2) (Table 1). IP_3_R2, mGluR5, PLC*β*, DAGL*α*, PLD2, and COX1 were found enriched in endfeet in at least one database while PP, MAGL, and PGES3 were expressed in the entire astrocyte with not overexpression in endfeet, and PLC*δ* was statistically more expressed in other compartments than in endfeet.

**Table 1.**
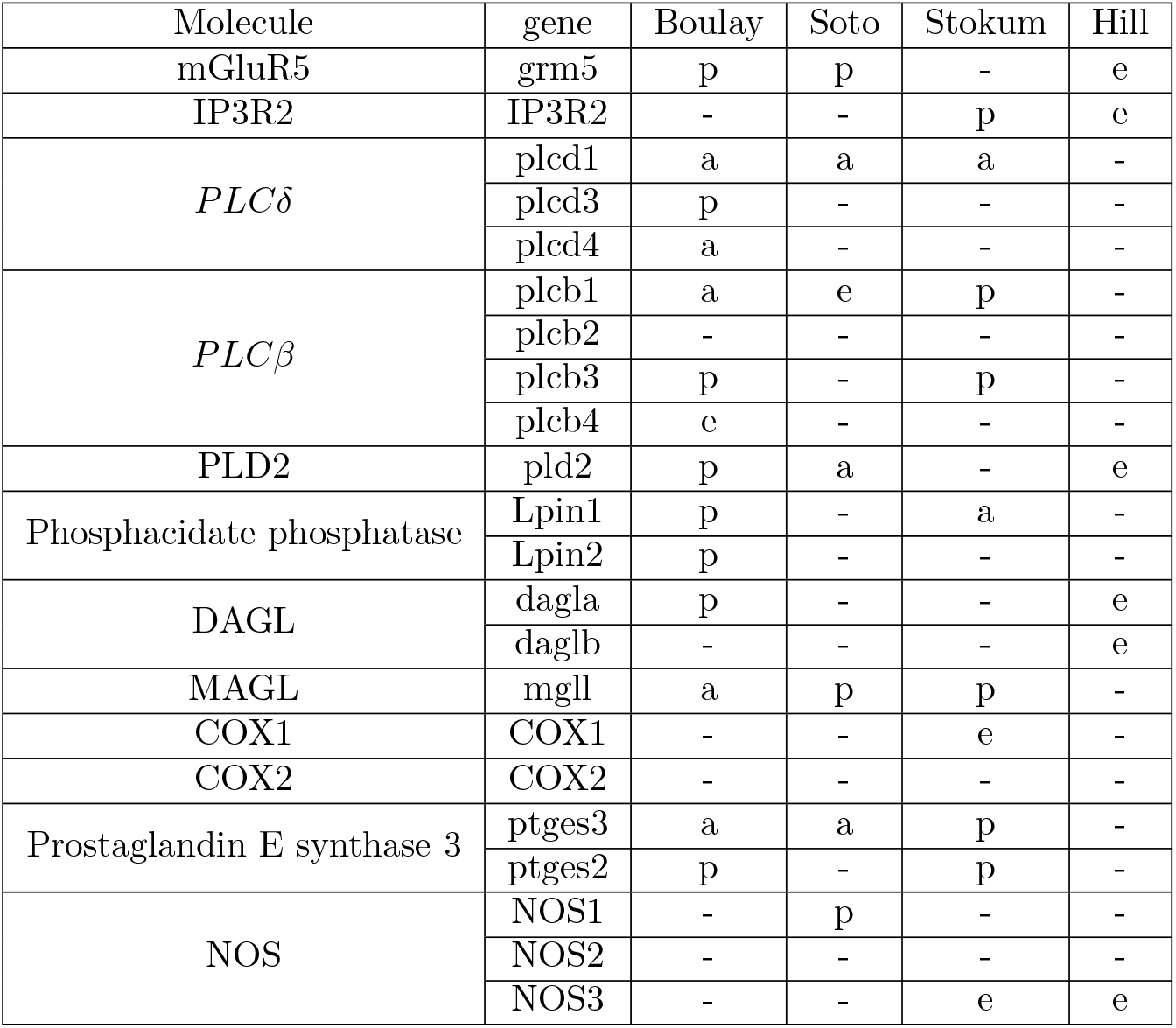
Summary of the expression of genes or proteins of the PGE2-pathway in four astrocyte omics datasets: Boulay et al. [29] (ribosome-bound mRNAs), Soto et al. [32] (proteomics), Stokum et al. [31] (proteomics), and Hill et al. [30] (proteomics). - : not found, e : found and statistically more expressed in endfeet, a : found and statistically more expressed in non-endfoot compartments, p : found, with no preferential location.

| Molecule | gene | Boulay | Soto | Stokum | Hill |
| --- | --- | --- | --- | --- | --- |
| mGluR5 | grm5 | p | p | - | e |
| IP3R2 | IP3R2 | - | - | p | e |
| *PLC*δ | plcd1 | a | a | a | - |
|  | plcd3 | p | - | - | - |
|  | plcd4 | a | - | - | - |
| *PLC*β | plcb1 | a | e | p | - |
|  | plcb2 | - | - | - | - |
|  | plcb3 | p | - | p | - |
|  | plcb4 | e | - | - | - |
| PLD2 | pld2 | p | a | - | e |
| Phosphatidate phosphatase | Lpin1 | p | - | a | - |
|  | Lpin2 | p | - | - | - |
| DAGL | dagla | p | - | - | e |
|  | daglb | - | - | - | e |
| MAGL | mgll | a | p | p | - |
| COX1 | COX1 | - | - | e | - |
| COX2 | COX2 | - | - | - | - |
| Prostaglandin E synthase 3 | ptges3 | a | a | p | - |
|  | ptges2 | p | - | p | - |
| NOS | NOS1 | - | p | - | - |
|  | NOS2 | - | - | - | - |
|  | NOS3 | - | - | e | e |

To facilitate the exploration of these datasets, we provide a script that queries and aggregates the data from these four studies. The code is available here.

From these expression data, we identify two main sources of PGE2 synthesis in astrocytes: phosphatidylcholine (PC) and phosphatidylinositol 4,5-bisphosphate (PIP2) [7, 38–45]. PC is hydrolyzed by phospholipase D2 (PLD2) into phosphatidic acid (PA) [38]. Both PA [39] and PIP2 [40] are substrates for diacylglycerol (DAG) production, which leads to PGE2 production by successive enzymatic reactions, which we will refer to as the PGE2 cascade. PA and PIP2 degradation into DAG are respectively catalyzed by phosphatidate phosphatase (PP) and phospholipase C *β* (PLC*β*) or *δ* (PLC*δ*). PLC*β* is activated by glutamate binding to mGluR while PLD2 is activated by Ca^2+^ [46]. PGE2 cascade then occurs according to the following steps:

DAG is converted into 2-Arachidonoylglycerol (2-AG) by diacylglycerol lipase (DAGL) [44], followed by arachidonic acid (AA) production by monoacylglycerol lipase (MAGL) [45], prostaglandin *H*_2_ (PGH2) synthesis by COX1 or COX2 [41], and PGE2 production by prostaglandin E synthase 3 (PGES3) [42]. The reaction scheme is depicted in Fig 1. We model it here using the set of differential equations described in the next subsections.

### Astrocyte Ca^2+^ and PGE2 signaling model

#### Calcium-induced calcium release model

Since PGE2 production is partly Ca^2+^ -dependent via PLD2 activation and because mGluR activation triggers calcium-induced calcium release (CICR), we included CICR in the model, using the G-ChI model proposed in refs [33, 47]. The G-ChI model is an extension of the Li-Rinzel model [48] that takes into account the dynamics of [*IP*3], and its Ca^2+^ -dependence. It describes the dynamics of the following variables: astrocyte cytosolic calcium concentration ([*Ca*^2+^]), IP_3_ concentration ([*IP*3]), IP_3_R2 opening probability (*h*), DAG concentration ([*DAG*]), classical protein kinase C concentration ([*cPKC*]), fraction of activated mGluRs (Γ), and extracellular glutamate concentration ([*G*]). The equations and parameter values of this model can be found in supplementary materials S1 and table S1, respectively. Representative temporal traces of [*Ca*^2+^], *h*, and [*IP*_3_] are shown in figure 2C.

We introduce here an extension of the model, called “PG-ChI”, which additionally takes into account the contribution of PA to DAG production:

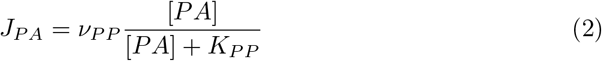

PLD2 activity, catalyzing PA synthesis from PC, is Ca^2+^ -dependent [46, 49]. To account for this regulation, we assume that Ca^2+^ modulates PLD2 activity in a saturable manner, consistent with enzyme activation kinetics. We therefore describe PA production by PLD2 using the Hill equation:

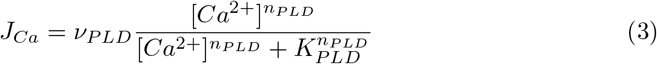

The mean field dynamics of PA concentration is thus given by:

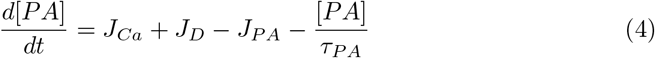

where *J*_*D*_ is the DAGK-mediated PA production rate taken here from the G-ChI model [33].

In the G-ChI model, DAG degradation is described by the *J*_*A*_ term, that takes into account DAG degradation into 2-AG by DAGL and other unknown DAG degradation mechanisms, the latter being assumed to be linear by the authors [33]. Here, we introduce an explicit *J*_2*AG*_ flux, which corresponds to the rate of DAGL-mediated production. Experimental reports indicated that 2-AG production by DAGL has a Ca^2+^ -dependent and a Ca^2+^ -independent component [43, 44, 50]. We thus combine a basal and a Ca^2+^ -dependent term:

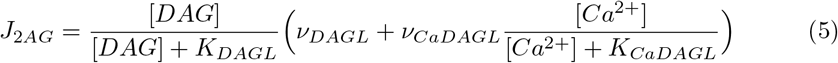

The mean field dynamics of DAG concentration thus becomes:

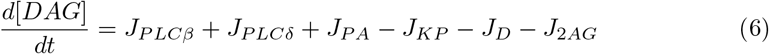

where *J*_*PLCβ*_ and *J*_*PLCδ*_ are the DAG production terms by PLC *β* and *δ*, respectively, and *J*_*KP*_ is the cPKC production term [33].

#### PGE2 cascade

The PGE2 cascade is modeled here with the following ODEs:

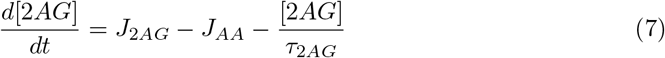

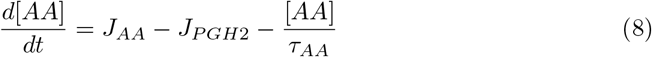

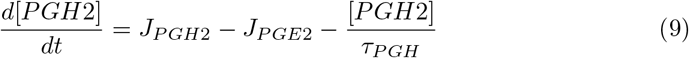

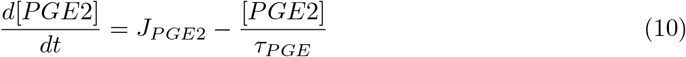

where:

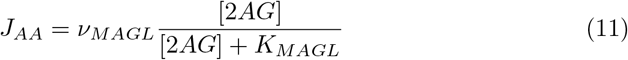

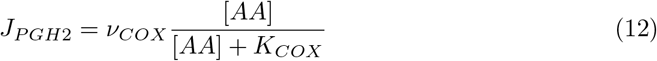

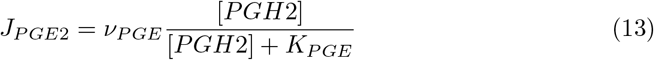

We could not find quantitative experimental data for MAGL kinetics. We therefore assume that MAGL-mediated 2-AG hydrolysis operates much faster than DAGL-mediated production. Thus, 2-AG dynamics is considered at quasi-steady state and 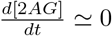. Neglecting 2AG degradation, i.e. 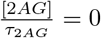, we obtain J_2AG_ ≃ J_AA_. Thus, the mean field dynamics of AA concentration, eq.(8) above, becomes:

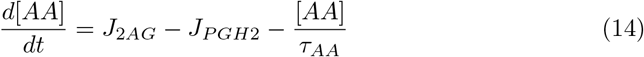

The combined PG-ChI and PGE2 cascade equations, eqs. (2)-(7) and (9)-(14), are referred to as the PG-ChI–PGE2 model. Parameter values of the PGE2 cascade model are listed in table 2 and parameter values of the G-ChI model in Supplementary table S1. The fluxes of the astrocyte Ca^2+^ and PGE2 signaling model are depicted in supplementary figure S3.

**Table 2.** Parameter values of the PGE2 cascade model.

| Parameter | Description | Value | Unit | Reference |
| --- | --- | --- | --- | --- |
| **PLD2 activity** |  |  |  |  |
| $\nu_{\text{PLD}}$ | Maximal rate of PA production | 1.2 | $\mu\text{M.s}^{-1}$ | - |
| $K_{\text{PLD}}$ | $\text{Ca}^{2+}$ affinity to PLD2 | 0.2 | $\mu\text{M}$ | - |
| $n_{\text{PLD}}$ | Hill coefficient of PA production | 2 | - | - |
| **PP activity** |  |  |  |  |
| $\nu_{\text{PP}}$ | Maximal rate of DAG production from PA | 5 | $\mu\text{M.s}^{-1}$ | - |
| $K_{\text{PP}}$ | PA affinity to PP | 12 | $\mu\text{M}$ | [39] |
| **DAGL activity** |  |  |  |  |
| $\nu_{\text{DAGL}}$ | Maximal rate of $\text{Ca}^{2+}$ -independent AA production | 0.055 | $\mu\text{M.s}^{-1}$ | - |
| $K_{\text{DAGL}}$ | DAG affinity to DAGL | 75 | $\mu\text{M}$ | [44] |
| $\nu_{\text{CaDAGL}}$ | Maximal rate of $\text{Ca}^{2+}$ -dependent AA production | 1 | $\mu\text{M.s}^{-1}$ | - |
| $K_{\text{CaDAGL}}$ | $\text{Ca}^{2+}$ affinity to DAGL | 2.4 | $\mu\text{M}$ | - |
| **COX1 activity** |  |  |  |  |
| $\nu_{\text{COX}}$ | Maximal rate of PGH2 production | 1 | $\mu\text{M.s}^{-1}$ | - |
| $K_{\text{COX}}$ | AA affinity to COX1 | 10 | $\mu\text{M}$ | [41] |
| **PGES activity** |  |  |  |  |
| $\nu_{\text{PGE}}$ | Maximal rate of PGE2 production | 1.5 | $\mu\text{M.s}^{-1}$ | - |
| $K_{\text{PGE}}$ | PGH2 affinity to PGES3 | 14 | $\mu\text{M}$ | [42] |
| **cAMP production** |  |  |  |  |
| $\nu_{\text{cAMP}}$ | Maximal rate of cAMP production | 2 | $\mu\text{M.s}^{-1}$ | - |
| $\text{EC}_{50}$ | PGE2 affinity to $EP_4$ receptors | 0.0003 | $\mu\text{M}$ | [51] |
| **Decay times** |  |  |  |  |
| $\tau_{\text{PA}}$ | PA decay time | 0.5 | s | - |
| $\tau_{\text{AA}}$ | AA decay time | 2 | s | - |
| $\tau_{\text{PGH}}$ | PGH2 decay time | disregarded | s | - |
| $\tau_{\text{PGE}}$ | PGE2 decay time | 1 | s | - |
| $\tau_{\text{cAMP}}$ | cAMP decay time | 1 | s | - |

### Vasculature model

To model the impact of PGE2 release on blood vessel diameter, a vasculature compartment was added to the endfeet compartment described above. The model is an extension of the three-compartment Windkessel model, developed by previous computational studies [28, 34, 35, 52]. It consists of an electrical circuit analogy that describes the mean field dynamics of the relative volumes of arterioles (*V*_*a*_), capillaries (*V*_*c*_), and venules (*V*_*v*_):

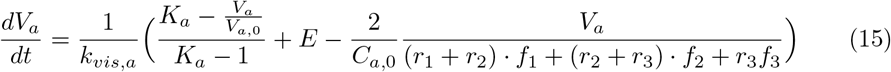

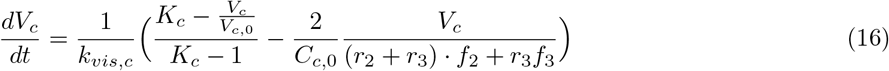

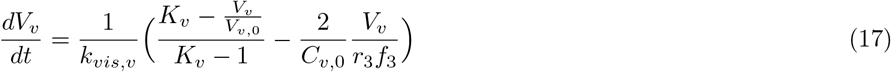

Assuming all blood vessels are cylinders of constant height, their relative diameter dynamics is given by:

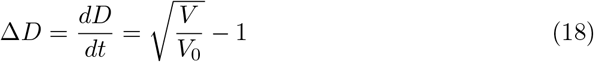

where *V* is the relative vessel volume and *V*_0_ the basal value of the relative vessel volume. Representative temporal traces of arterioles, capillaries, and venules relative diameters are shown in figure 2D.

The influence of external factors on blood vessel radius is described by *E*, which is defined in the next subsection. Parameter values of the vasculature model were taken from [28] and are displayed in supplementary table S2.

### Coupling of neuron, astrocyte, and vascular dynamics

Arteriole smooth muscle cells express four G protein-coupled receptor subtypes that are activated by PGE2: *EP*_1−4_. *EP*_2_ and *EP*_4_ activation triggers vasodilation [53, 54]. *EP*_4_ is weakly expressed in cerebral arteriolar smooth muscle cells (SMCs) [55, 56], whereas *EP*_2_ does not seem to be expressed. Similar observations were made in human middle cerebral arteries [54]. We therefore restrict the focus of our model to *EP*_4_.

*EP*_4_ activation by PGE2 results in an increase of *cAMP* concentration in the SMC [51, 57], so that the mean field dynamics of cAMP concentration can be described as:

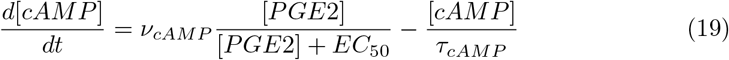

cAMP then inhibits Myosin-Light-Chain phosphatase (MLCP) via PKA stimulation, which in turn phosphorylates MLCP, resulting in cytosolic Ca^2+^ decrease and subsequent vessel dilation [58–60]. Since this mechanism is not fully understood we model the influence of cAMP concentration in vascular SMC [*cAMP*] on arteriole diameter as a Hill function with exponent *n*_*c*_. In addition to cAMP, the arteriole diameter is also directly impacted by neuronal NO production. The influence of neuronal and astrocyte activity on arteriole diameter (term *E* in eq.(15) above) is therefore modeled as:

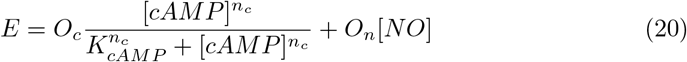

where *O*_*c*_ is the maximal cAMP effect on arteriole dilation, *K*_*cAMP*_ is the half cAMP activation concentration, and *O*_*n*_ is the maximal NO effect on arteriole dilation.

### Spatial model

To assess the impact of the location of neuronal stimulation on astrocyte-mediated NVC, we spatialized the astrocyte compartment using the finite volumes method [61]. Astrocyte geometry was simplified into three compartments: a soma, an endfoot, and two leaflets connected by branches (Figure 6A). Branches are themselves made of cylinders sharing the same diameter d. Each portion of a branch connecting two compartments (endfoot, soma, leaflets), or a compartment and an intersection between two branches, is divided into ten cylindrical subcompartments.

The endfoot volume was set to 2.0 *µ*m^3^ based on unpublished 3D endfoot reconstructions from Focused Ion Beam-Scanning Electron Microscopy (FIB-SEM) data (courtesy from Prof. K. Murai, McGill University, Montreal, Canada). The diameter of the astrocytic soma was estimated based on a confocal image of an astrocyte expressing GCaMP6f from Covelo et al. (ref [62], Fig.1). Since the soma was modeled as a sphere, its volume was set to 4200 *µ*m^3^. Lastly, the volume of the leaflets was set to 0.4 *µ*m^3^, based on FIB-SEM data of leaflet perisynaptic astrocytic processes [63, 64]. These parameter values are listed in table 3. The values of branches diameter d and lengths L_1_ and L_2_ varied in our simulations to reflect their experimentally reported variability: from 10-200 nm (branchlets) to 1-2 *µ*m (branches) in diameter and up to 20 *µ*m in length [62, 65] (table 4).

**Table 3.** Constant geometrical parameter values of the spatial model. Length is the compartment size along the diffusion axis. The whole astrocyte geometry is displayed in Fig 6.

| Compartment | Volume ($\mu\text{m}^3$) | Length ($\mu\text{m}$) |
| --- | --- | --- |
| Soma | 4200 | 20 |
| Endfoot | 2 | 1 |
| Leaflets | 0.4 | 0.1 |

**Table 4.** Varying geometrical parameter values of the spatial model. These parameters describe the sizes of branches connecting the somatic, endfoot, and leaflet compartments. Branches are modeled as cylinders. The whole astrocyte geometry is displayed in Fig 6.

| Parameters | $L_1$ | $L_2$ | d |
| --- | --- | --- | --- |
| Variation interval ($\mu\text{m}$) | 1-20 | 1-15 | 0.02-2 |

All subcompartments (endfoot, soma, leaflets, and cylinders) are coupled via one-dimensional Ca^2+^ and IP_3_ diffusion. According to Fick’s first law, the molar flux density *j* is proportional to the concentration gradient ∇*C*:

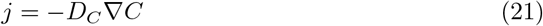

where *D*_*C*_ is the diffusion coefficient.

Assuming one-dimensional diffusion along the x-axis, the equation becomes:

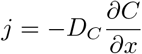

Let *S* denote the exchange surface between two neighboring compartments. The total molar flux across this surface is then given by:

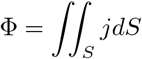

Considering a uniform flux density over the surface, this expression simplifies to:

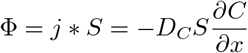

The mean-field dynamics of the molecule concentration in compartment i *C*_*i*_, which is coupled by diffusion to compartments *i* + 1 and *i* − 1 is thus given by:

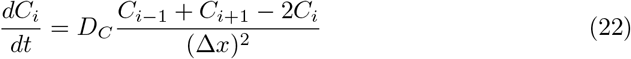

where Δ*x* is the distance between neighboring compartments. See supplementary materials S2 for further details.

Assuming that DAGL is only expressed in endfeet (see table 1), the set of reactions occurring in the endfoot compartment was described by the PG-ChI–PGE2 model, whereas the PG-ChI model was implemented in the other compartments, including branches.

### Simulations

We used the solve ivp function from the scipy.integrate library, with the Radau algorithm, to simulate the dynamics of the well-mixed system (18 ODEs) and the spatialized model (282 ODEs). The code, implemented in Python, is available here. Parameter values were taken, whenever possible, from the literature. The remaining parameters were calibrated to fit experimental arteriole Δ*D* traces, within biologically plausible ranges: maximal rate of molecules production were chosen between 0.01 and 10 *µM*.*s*^−1^; decay times were taken between 0.1 and 5 s, or disregarded, whenever possible, for model simplicity. Model dynamics was computed at rest conditions (no neuronal stimulation), until the equilibrium was reached (≈ 50 s simulation time). The steady state values were used as the initial conditions of all the simulations presented in this study.

### Data analysis

The Root Mean Squared Error (RMSE) was computed using numpy to quantify the distance between virtual and experimental Δ*D*. The bifurcation analysis was computed with the XPPAUT software. For computational simplicity, [*G*] was treated as a constant parameter for the bifurcation analysis (bifurcation parameter), instead of the pulsatile input of the full model.

## Results

### The model successfully reproduces *in vivo* arteriole dilation dynamics

To study the role of astrocytes in NVC, we have developed an ODE model that couples glutamatergic neuronal activity, astrocyte activity, and blood vessel diameter variations. The reaction scheme of this model is presented in Figure 1. The model of astrocyte activity was chosen based on astrocyte translatome (ribosome-bound mRNAs, [29]) and proteomics datasets [30–32] and couples the Ca^2+^ and PGE2 signaling pathways. Astrocyte Ca^2+^ activity was implemented based on the G-ChI model [33, 47], which simulates Ca^2+^ -induced Ca^2+^ release. The extension proposed here, referred to as the PG-ChI-PGE2 model, additionally accounts for the PGE2 signaling pathway as well as PA-mediated DAG production. Neuronal activity is modeled as a release of glutamate and NO, as the latter has been reported to partly mediate NVC [66, 67]. An extended three-compartments Windkessel model was implemented to simulate the dynamics of blood vessel diameter change, as done in previous computational studies [28, 34, 35, 52]. The model is implemented as a set of eighteen ordinary differential equations (see Methods section). Parameter values were taken from the literature when available and manually set otherwise (see table 2, and supplementary materials S1 and S2).

To validate the model, we first compared its dynamics with *in vivo* mice arteriole diameter recordings from Institoris et al. [8] (Figure 3). To investigate the role of astrocytes in NVC, Institoris et al. [8] used Adeno-Associated Viruses expressing a plasma membrane Ca^2+^ ATPase targeting astrocytes (CalEx), which reduces evoked and spontaneous elevations of free Ca^2+^ in astrocytes [68]. In the model, CalEx conditions were simulated by setting *d*[*Ca*^2+^]*/dt* = 0. The simulated relative arteriole radius variations, Δ*D* closely matched the experimental traces in control conditions under sustained neuronal stimulation (30 s), with a root mean squared error (RMSE) of 2.5 (Figure 3B). For short neuronal stimulations (5 s), the simulated traces qualitatively reproduced the experimental data, but the fit displayed a larger RMSE of 5.4 (Figure 3 A). The model was able to capture both the “early-phase” (from stimulation onset to ≈ 2-4 s post-stimulation) and the “late-phase” of NVC (from 5-7 s post-stimulation onset to stimulation termination). Interestingly, the model was also able to reproduce Δ*D* dynamics with reduced astrocyte Ca^2+^ activity (CalEx) under either short (Figure 3C, RMSE=4.0) or long (Figure 3D, RMSE=3.38) neuronal stimulation. Both simulation and experimental CalEx traces were characterized by a decreased blood vessel radius dilation compared to control conditions during the late-phase of NVC. Note that vessel dilation was delayed by a few seconds in the CalEx simulations under short stimulation duration, which explains the increased RMSE between simulated and experimental data in these conditions. Importantly, the simulated radius dynamics following short neuronal stimulation displays a secondary radius dilation peak that is delayed compared to experimental traces (Figure 3A). As it is virtually absent from the simulated CalEx trace (Figure 3C), the second peak is likely mediated by astrocyte Ca^2+^ signaling.

**Fig 3.**
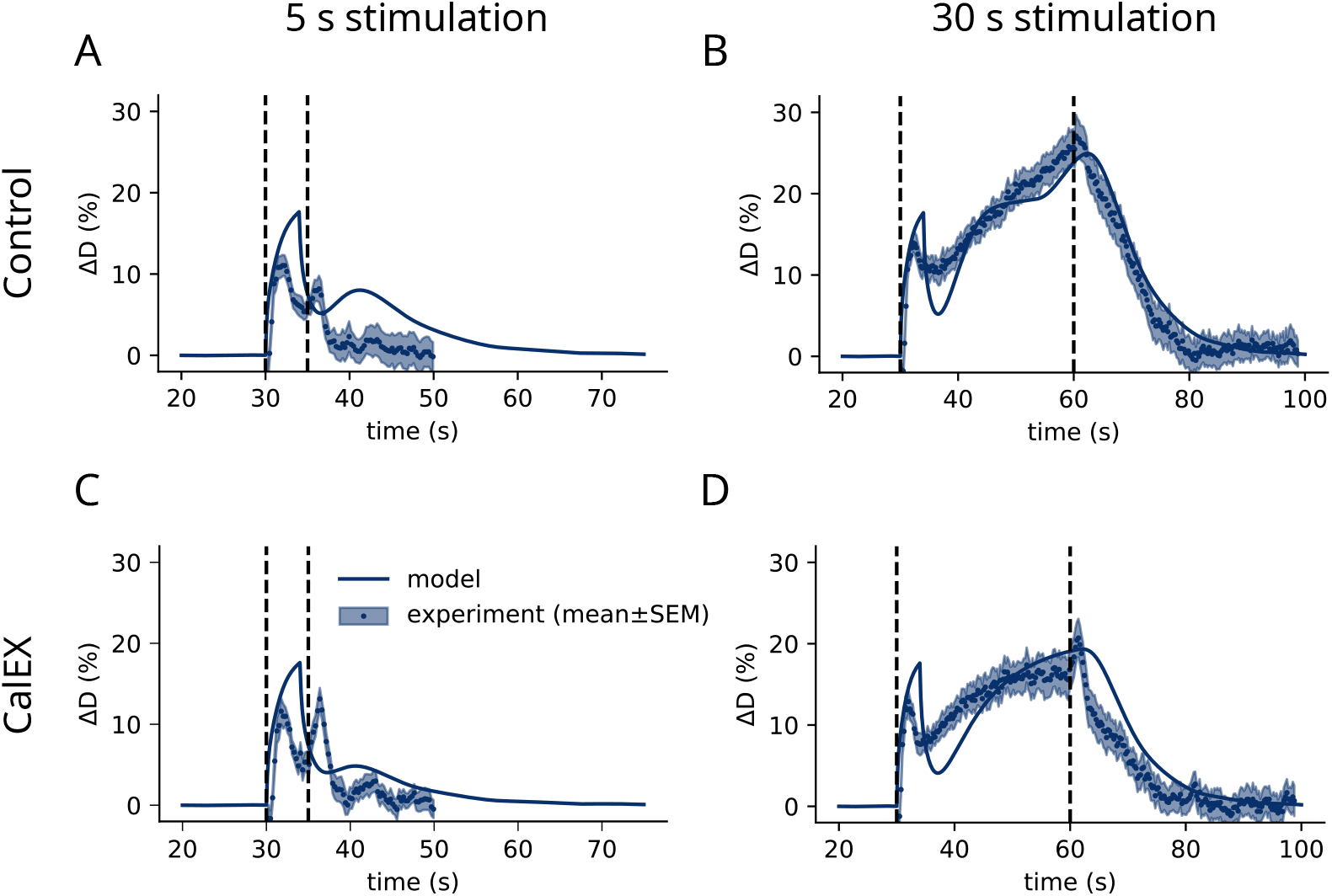
The model successfully reproduces *in vivo* experimental arteriole diameter dynamics. Temporal variations of the simulated relative arteriole radius (dark blue line), Δ*D*, in control (top, A-B) and CalEX (bottom, C-D) conditions, under 5 s (left, A,C) and 30 s (right, B,D) neuronal stimulation. Experimental traces from Institoris et al. [8] are displayed as mean values (blue dots) *±*SEM. Black vertical dashed lines delimit the neuronal stimulation period.

### The PGE2 pathway triggers the late phase of NVC

In order to investigate the contribution of the astrocyte PGE2 and neuronal NO pathways to NVC, we ablated each pathway from the model independently, yielding the “PGE2-KO” and “NO-KO” models, respectively. While the NO pathway allowed for rapid diameter dilation displaying similar characteristics to the “early-phase” of NVC, the PGE2 pathway triggered a slower and delayed dilation reminiscent of the “late-phase” of NVC, both in control and CalEx conditions (Figure 4A). To gain a better understanding of the contribution of each element of the PGE2 pathway to NVC, we knocked out specific enzymes of the signaling cascade by setting their maximum production rate value to 0 (Figure 4B). Notably, inhibiting PLD2 activity (*ν*_*P LD*_=0 in equation 3) resulted in a decrease of the maximum value of Δ*D*. Note that knocking down PP (*ν*_*P P*_ = 0 in equation 2) had similar effects (fused curves; not shown).

**Fig 4.**
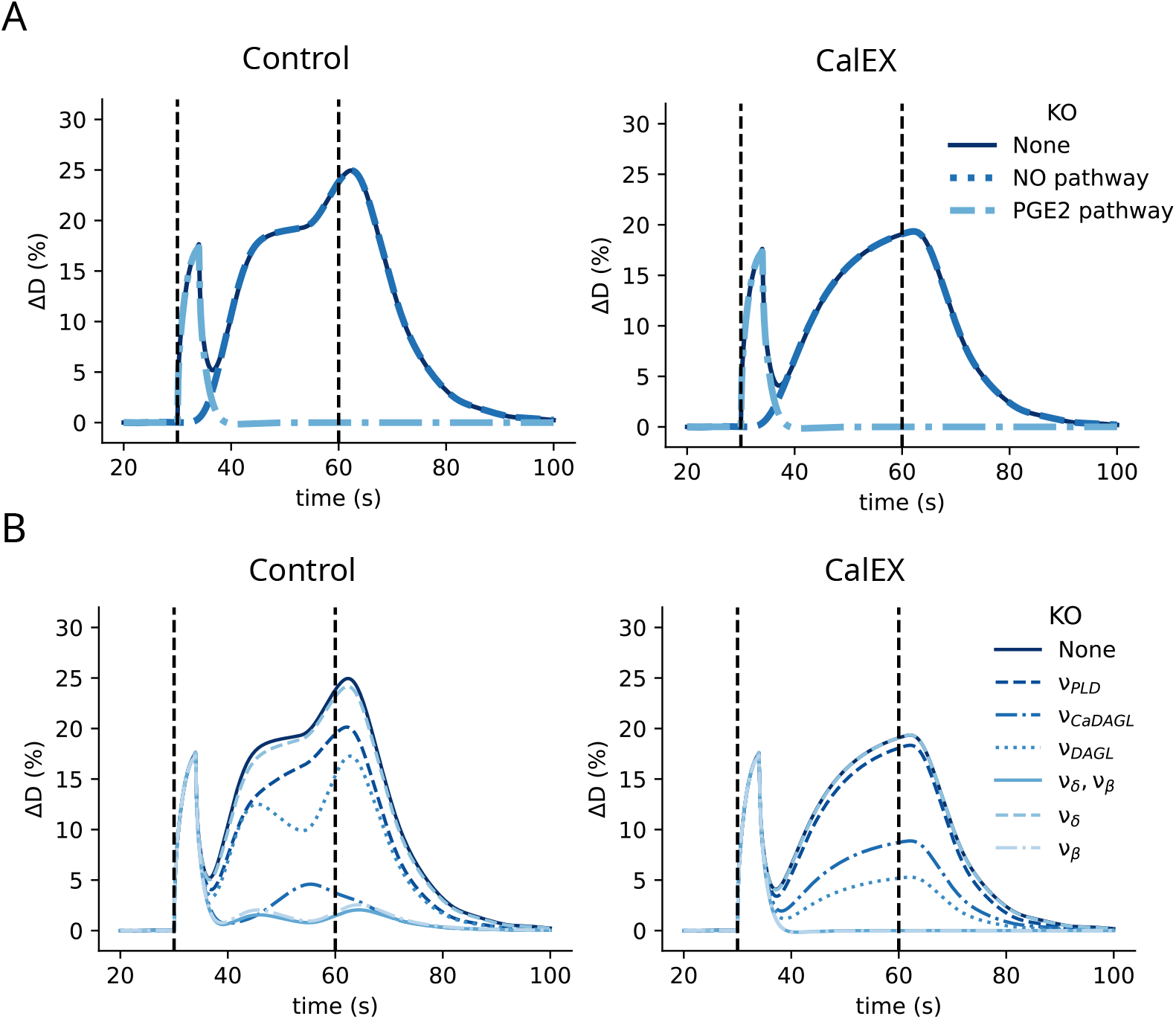
Study of the contribution of modeled reactions to the different phases of NVC. (A) Comparison of the temporal variations of the relative arteriole radius, Δ*D*, in the control (dark blue line), NO-KO (blue dotted line) and PGE2-KO (light blue dash-dotted line) models in control (left) and CalEX (right) conditions. (B) Temporal variations of Δ*D* for another set of virtual KOs: *ν*_*P LD*_-KO, *ν*_*DAGL*_-KO, *ν*_*CaDAGL*_-KO, *ν*_*δ*_-KO, and *ν*_*β*_-KO models in control (left) and CalEX (right) conditions. Black vertical dashed lines delimit the neuronal stimulation period.

Then, we studied the influence of *PLCβ*- and *PLCδ*-mediated DAG production on NVC. First, we knocked out both PLC isoforms, (*ν*_*δ*_ = 0 and *ν*_*β*_ = 0). To specifically evaluate the role of DAG on NVC, parameter values were only modified for DAG production, whereas those for IP_3_ production were kept unchanged. Surprisingly, most of the diameter dilation was abolished in the “late-phase”, with Δ*D* reaching lower values than when knocking down PLD2. In this model, *PLCβ* strongly influences NVC while *PLCδ* has minimal effects, as shown by the specific effect of *PLCβ* and *PLCδ* KO (Figure 4B). This is not surprising as *ν*_*β*_ is about ten times larger than *ν*_*δ*_ [33], which is consistent with the low expression of *PLCδ* in endfeet compared to the rest of the astrocyte (table 1). Note that suppressing *PLCδ*-mediated production of both DAG and IP_3_ was not significantly different from suppressing *PLCδ*-mediated DAG production alone (fused curves; not shown). In contrast, inhibiting both *PLCβ*-mediated IP_3_ and DAG production completely abolished the “late-phase” of NVC (not shown).

To evaluate the respective roles of the Ca^2+^ -dependent and Ca^2+^ -independent DAGL activity in NVC, we respectively set *ν*_*CaDAGL*_ and *ν*_*DAGL*_ to 0. Knocking down *ν*_*DAGL*_ resulted in a decreased maximum Δ*D* by roughly 30 %. The amplitude of vessel diameter oscillations was about 75 % higher than in control conditions, as in the latter Δ*D* oscillations are smoothed by the Ca^2+^ -independent DAGL effect. In contrast, suppressing the Ca^2+^ -dependent DAGL effect *ν*_*CaDAGL*_ altered Δ*D* oscillations by reducing the maximal amplitude of Δ*D* by 80 %.

### Increased extracellular neuronal glutamate release results in larger arteriole dilation and astrocyte Ca^2+^ oscillations

As our results suggest that glutamate is causal to the “late-phase” of NVC (Figure 4, we hypothesized that the amount of neuronal glutamate released at the vicinity of the astrocyte should strongly impact astrocyte activity and blood vessel dilation. To test this hypothesis, we ran simulations with varying values of *ν*_*G*_, the peak glutamate concentration occurring at each stimulation time *t*_*spike*_ (equation 1). As expected, decreasing *ν*_*G*_ did not affect the early-phase of NVC but resulted in a reduced “late-phase” maximum relative arteriole diameter. For example, a 80 % decrease in *ν*_*G*_ resulted in a 60 % decrease in the maximum “late-phase” Δ*D* (Figure 5A). Moreover, a tenfold decrease in glutamate amplitude completely supressed the second peak of Δ*D*. A bifurcation analysis of the system reveals that a subcritical Hopf bifurcation occurs when the extracellular glutamate concentration [*G*] reaches about 2.4 *µM* (Figure 5B). Above this value, [*Ca*^2+^], [*IP*_3_] and [*DAG*] oscillate, leading to the observed Δ*D* oscillations.

**Fig 5.**
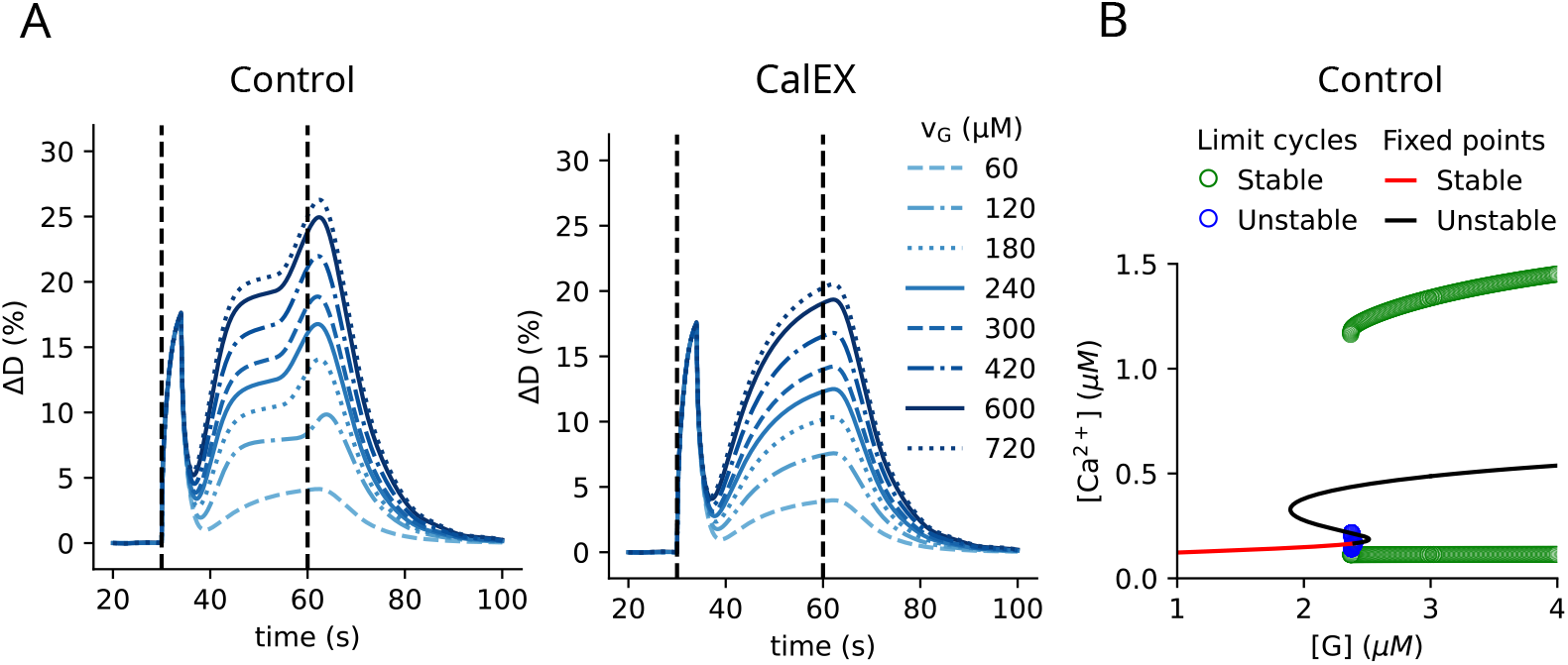
Extracellular glutamate concentration governs arteriole dilation and astrocyte Ca^2+^ oscillations. (A) The peak extracellular glutamate concentration, *ν*_*G*_, strongly impacts the relative blood vessel diameter variations, Δ*D*, following neurostimulation in control (left) and CalEX (right) conditions. Δ*D* oscillations can be observed from *ν*_*G*_ = 120 *µM*, in control conditions only. Black vertical dashed lines delimit the stimulation period. (B) Bifurcation diagram of the model with the extracellular glutamate concentration [*G*] as a bifurcation parameter. A subcritical bifurcation appears for [*G*] ≈ 2.4 *µ*M, giving rise to an unstable (blue circles) and a stable limit cycle (green circles). The unstable fixed point (black line) resulting from the Hopf bifurcation however coexists with two other unstable fixed points. The unstable limit circle collides with the intermediate unstable fixed point close to the Hopf bifurcation.

### Peri-endfoot glutamatergic release enhances astrocyte PGE2-mediated NVC

Different astrocyte compartments, including endfeet, have recently been reported to contact synapses by Aten et al. (ref. [69] Fig. 6). Therefore, we next investigated the impact of the location of active synapses on astrocyte-mediated NVC. To do so, we implemented a spatial model of astrocyte-dependent NVC. The astrocyte in our spatial model is composed of the following compartments: a soma (S), an endfoot (Ef), leaflets (F_1_ and F_2_), and branches of diameter d and lengths L_1_ and L_2_, coupled by Ca^2+^ and IP_3_ diffusion (Figure 6A). DAG, a membrane lipid, was not considered a diffusing molecule [70, 71]. In accordance with recent astrocyte omics data suggesting that DAGL and COX1 are only expressed in endfeet (table 1), the PGE2 signaling cascade was only implemented in the endfoot (PG-ChI–PGE2 model, see Methods section for details). To explore the impact of astrocytes on NVC depending on the nature and spatial distribution of the stimulated compartment, simulations were run for different values of d, L_1_, and L_2_, following either somatic (S), endfoot (Ef), or leaflet (F_1_ or F_2_) stimulation. Note that since astrocytes only impact the “late-phase” of NVC (Figure 4), only the latter was analyzed in this set of *in silico* experiments.

**Fig 6.**
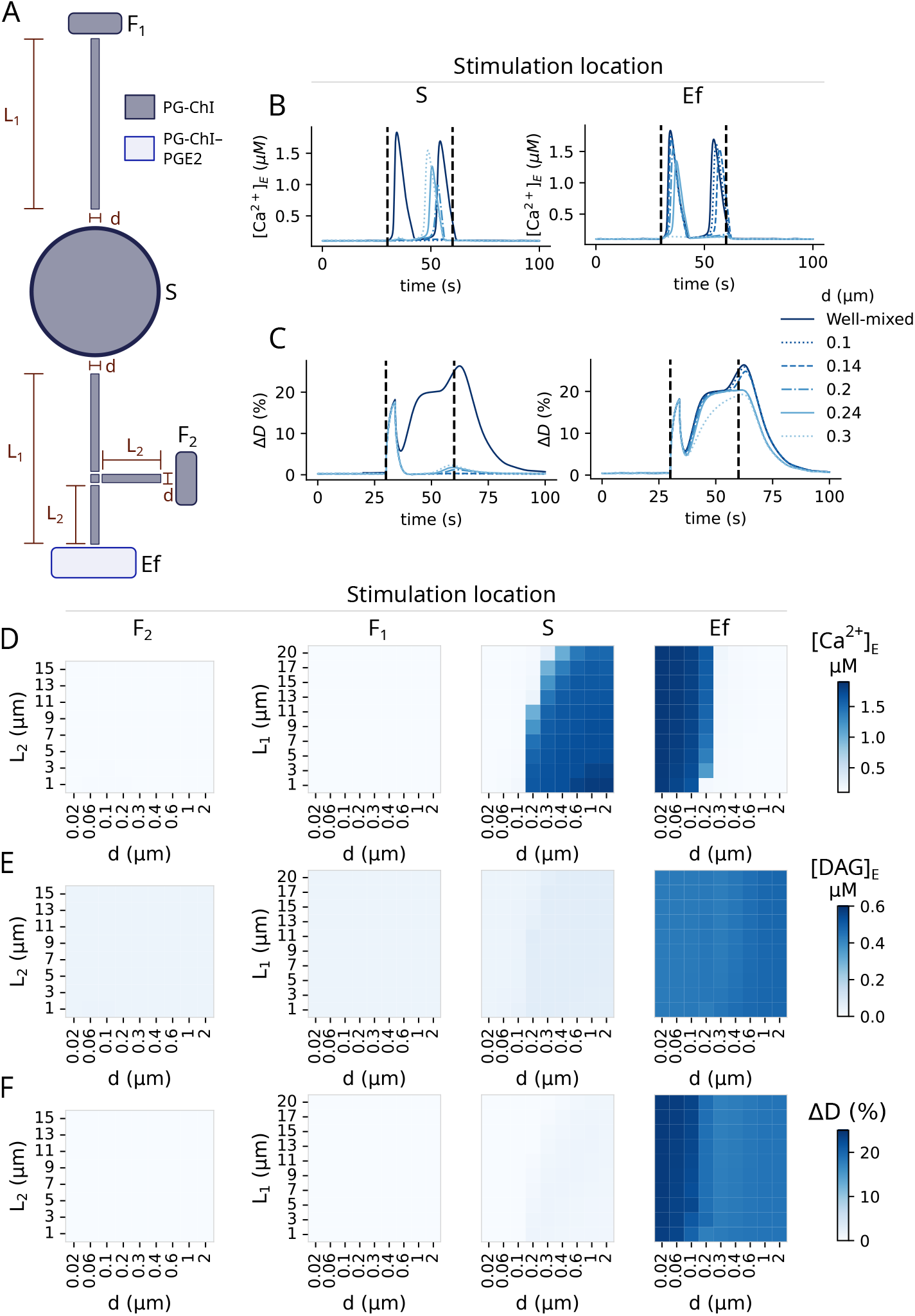
Simulations of the spatial model suggest that astrocyte PGE2-mediated vasodilation only happens when neuronal stimulation occurs at the endfoot level. (A) Schematic representation of the simplified astrocyte geometry in the spatial model. It is composed of a soma (S), an endfoot (Ef), leaflets (F_1_ and F_2_), and branches of diameter d and lengths L_1_ and L_2_. Traces of endfoot Ca^2+^ concentration (B) and Δ*D* (C) for a range of branch diameter d in simulations where either the soma (S, left), or the endfoot (E, right) was stimulated, L_1_ = 11 *µ*m and L_2_ = L_1_*/*2. Black vertical dashed lines delimit the neuronal stimulation period. Heatmaps show the peak endfoot concentration of Ca^2+^ ([Ca^2+^]_E_, D) and DAG ([DAG]_E_, E), and the peak relative arteriole dilation (Δ*D*, F), for different stimulation locations: leaflets (F2 or F1), soma (S), or endfoot (Ef). In *in silico* experiments, L_1_ was set to 20 *µ*m when varying L_2_. L_2_ was set to L_1_*/*2 when varying L_1_.

First, whatever the size of the compartments and parameter values, Ca^2+^ signals were never found to reach the endfoot when neuronal stimulation occurred at leaflets (fig.6D/F_1_,F_2_). Only soma or endfoot stimulation yielded Ca^2+^ signals that were transmitted to the endfoot (fig.6D/S,Ef). In this case, the influence of branch diameter d on the amplitude of the transmitted Ca^2+^ signal was completely different depending on whether neuronal stimulation occurred at the soma or at the endfoot levels. The amplitude of the endfoot Ca^2+^ signal increased with d for somatic stimulation but decreased with increasing d under endfoot stimulation. Indeed, large values of d facilitate the diffusion of Ca^2+^ from the soma to the endfoot for somatic stimulation, but they facilitate Ca^2+^ diffusion away from the endfoot upon endfoot stimulation. Therefore, our spatial model suggests that endfoot Ca^2+^ signals are favored by large branch diameters if neuronal stimulation happens on the soma whereas they are favored by small branch diameters if neuronal stimulation occurs directly at the endfoot. In contrast, endfoot DAG concentration was barely affected by astrocyte geometry, probably because it does not diffuse.

In strong opposition to Ca^2+^ dynamics, however, endfoot DAG production and blood vessel dilation mainly occurred when the stimulated compartment was the endfoot and not the soma. The smaller the astrocyte branch diameter, the larger the arteriole dilation. More precisely, endfoot stimulation in astrocyte geometries with branch diameter *d <* 0.1 *µ*m displayed relative arteriole diameter increases Δ*D* similar to the well-mixed model (Figure 6C). Above this value, Δ*D* decreased to about 15 %. Simulations with values of *d >* 0.2 *µ*m suggest that endfoot DAG concentration increases are sufficient to trigger arteriole dilation even in the absence of Ca^2+^ signals (Figure 6E-F). We also note that somatic stimulation can trigger Ca^2+^ signals in the endfoot that reach levels similar to the well-mixed model, but these somatically-triggered Ca^2+^ signals resulted in minor peaks of endfoot DAG concentration and, consequently, in minor Δ*D* elevations (under 3 %). Moreover, somatic stimulation failed to trigger the first Ca^2+^ peak compared to the well-mixed implementation, but triggered the second Ca^2+^ peak up to 6 s earlier than the second peak of the well-mixed model (Figure 6B).

Because leaflets have recently been reported to form domains that integrate multiple synapses [72], we next simulated a domain of ten neighboring leaflets at either F_1_ or F_2_ locations (supplementary figure S1). Only F_2_ leaflets activation was able to trigger endfoot [Ca^2+^] elevations, and only when they were located very close to the endfoot (1 *µm*). Surprisingly, the influence of branch length on endfoot activity after endfoot or somatic stimulation was larger when multiple leaflets were active (compare supplementary figure S1 with figure 6). This phenomenon is particularly strong for branch diameters ≥200 *µ*m. This may result from an increase in the volume of the entire astrocyte with branch length, resulting in an enhanced dilution effect. As in the case of single leaflet activation (figure 6), the arteriole was mainly dilated when stimulation occurred at the endfoot level.

Lastly, we tested scenarios where the PGE2 pathway was expressed in all astrocyte compartments. To this end, we simulated the PG-ChI-PGE2 model in all modeled compartments (supplementary figure S2). As the diffusion coefficients of PA, AA, PGH2, and PGE2 were not available in the literature, they were assumed to equal 300 *µm*^2^.*s*^−1^. This value is an upward estimate, based on the effective Ca^2+^ and IP_3_ diffusion coefficients, respectively 13 and 280 *µm*^2^.*s*^−1^ [71]. Under these assumptions, the influence of the soma on NVC was larger than when the PGE2 pathway is restricted to the endfoot, triggering arteriole dilation to levels close to those of the well-mixed model. This effect was however only possible if the soma and the endfoot were connected by large branches (high value of d, figure S2). Conversely, the influence of endfoot activation on arteriolar dilation decreased more rapidly with d than when only endfeet expressed the PGE2 pathway molecules (figure S2 vs. 6). As more molecules are diffusing in this implementation of the model, the dilution effect was amplified.

Taken together, these results suggest that astrocyte geometry plays a significant role in shaping intracellular diffusion and therefore Ca^2+^ and PGE2 dynamics. Overall, neuronal stimulation at the leaflets level triggered almost no arteriole dilation, regardless of the geometric parameters. Neuronal stimulation at the somatic level could only trigger arteriole dilation if the soma was expressing molecules of the PGE2 pathway; if PA, AA, PGH2, and PGE2 diffused within the astrocyte; and if the diameter of branches connecting the soma to the endfoot was large. Neuronal stimulation at the endfoot level always triggered arteriole dilation, provided that only Ca^2+^ and IP_3_ were diffusing, with NVC being optimal when endfeet were connected to thin branches.

## Discussion

The roles of astrocytes in NVC remain highly debated. In particular, the involvement of astrocytic Ca^2+^ signals in NVC is controversial. Although Ca^2+^ signals have been observed in endfeet concomitantly with vasodilation [73–75], early studies suggested that these Ca^2+^ transients followed blood vessel dilation temporally, and therefore did not trigger it [14]. More recently, hyperemia has been observed *in vivo* in the absence of IP_3_R2-mediated Ca^2+^ signals in astrocytes [13]. In contrast, other *in vivo* experiments in awake mice challenged this view by suggesting that NVC could be partly astrocyte Ca^2+^ -mediated [8, 76]. Here, we have developed a model that accounts for these observations. The model recapitulates the astrocyte-independent early phase and partially astrocyte-dependent late phase of NVC reported by Institoris et al. [8]. The latter is triggered by PLD2 and Ca^2+^ -dependent DAGL activity, as well as Ca^2+^ -independent *PLCβ* and DAGL activity. Our model successfully replicates qualitative arteriole diameter variations both in control and CalEx conditions. Whether these two phases of NVC vary depending on blood vessel type, brain region, and other (patho-)physiological sources of variability of the neuro-gliovascular unit remains to be explored.

Neuronal NO is a key player of NVC [6, 67, 77]. It is produced upon N-methyl-D-aspartate (NMDA) receptor activation by the neuronal nitric oxide synthase isoform (nNOS). NO then diffuses and activates soluble guanylate cyclase in neighboring vessel smooth muscle cells, leading to cGMP synthesis, which downstream reactions result in SMC relaxation [5, 77]. Here, as we focus on astrocyte-mediated NVC, NO release is described with a simple phenomenological model. This implementation was enough to obtain good qualitative fitting of the model to *in vivo* experimental traces in awake mice. Interestingly, the relative contribution of NO to NVC varies with local expression levels, which are brain region-dependent [78]. A mechanistic model of neuronal NO signaling and downstream reactions in SMCs could help provide a better quantitative match of the model with experimental traces and take into account this local variability.

Our results suggest that blocking PLC*β* activity should result in a decrease in arteriole dilation during the late-phase of NVC. This is in contrast to experiments on brain slices that reported no reduction in stimulation-evoked capillary dilation following the pharmacological blockage of PLC [7]. This is not entirely surprising as mGluR blockage in this study did not affect endfoot Ca^2+^ activity either. The use of brain slices may have influenced this result since other studies on *in vivo* adult mice [79] or adult rats [74] have shown that astrocyte mGluR5 were partly mediating NVC. As the gliovascular unit varies depending on the developmental stage, brain region, and in disease, together with variations in cell and blood vessel composition, the signaling pathways involved are likely to vary too. For example, glutamatergic axons in the mouse cerebral cortex have been reported to dilate neighboring arterioles via direct synaptic-like neuron–arteriolar smooth muscle cell communication [80], and astrocytes have been suggested to contribute to NVC at the capillary but not at the arteriolar level [7]. The pathways involved in astrocyte release of vasodilators are also likely to vary. For example, astrocyte P2X purinoreceptors have been suggested to contribute to PGE2 release at the capillary level [7]. Lastly, depending on the nature of the neuro-gliovascular unit at stake, other cell types such as endothelial cells are likely to contribute to NVC [81–84]. Future *in vivo* experiments monitoring Ca^2+^ activity in endfeet and its dependence to PLC activity, as well as its reliance on the nature of the neighboring vascular structure, will be important to test and refine our understanding of the mechanisms dictating NVC.

In line with recent translatome [29] and proteomics data [30–32], the spatial implementation of the model assumes that PGE2 signaling is restricted to the endfeet. Indeed, mGluR and molecules of the PGE2 pathway are enriched in endfeet compared to other astrocyte sub-cellular compartments. Our *in silico* experiments suggest that stimulating the astrocyte elsewhere than the endfoot compartment would trigger attenuated or no endfoot Ca^2+^ elevations and hardly any arteriole dilation. In these simulations of our model, Ca^2+^ and IP_3_ could not diffuse quickly enough or in sufficient quantities from the stimulated compartments to the endfoot to trigger the same dilation as when the endfoot was stimulated. This result suggests that astrocyte-mediated NVC might occur locally, at the endfoot level. Interestingly, recent EM data support this view, as synapses have been observed in direct contact with endfeet [69]. As the number of synapses in contact with endfeet is likely to change depending on physiological conditions, we simulated the effect of the amount of glutamatergic input on blood vessel dilation. Our results suggest that an increased number of synapses contacting the endfoot would amplify the late-phase of NVC while a decrease in synaptic connectivity would impair NVC. Congruently, synapse loss and neurovascular uncoupling are observed in early stages of Alzheimer’s disease (AD), which contributes to cognitive decline [5, 10]. The temporality of neuro-gliovascular dysfunction in AD at the subcellular level remains to be explored and should provide novel insights into AD pathogenesis. Overall, our results, in line with previous reports [20], challenge the classical view that opposes perivascular and perisynaptic processes as different entities and suggest that endfeet could be both, which would allow them to act as modulators of NVC at the sub-cellular level. A thorough investigation of the synaptic environment of endfeet will be critical to refine the view of endfeet as perisynaptic entities. The development of computer vision tools tailored to astrocytes will be critical to achieving this goal [85].

The model presented here is a generic model of astrocyte-dependent neurovascular coupling, based on omics data capturing the ribosomal RNA expression in endfeet [29], the interactome of aquaporin 4 [32], a water channel enriched in endfeet, as well as the endfoot proteome [30, 31]. These datasets shed light on proteins expressed in a population of astrocytic endfeet but do not allow to discriminate proteins expressed by different astrocytes or different endfeet. Notably, most of these datasets are enriched in endfeet neighboring capillaries and might thus poorly reflect the proteome of endfeet at the arteriolar interface. Given the growing literature on astrocyte diversity in health and disease [86–89] and the local translation of proteins in endfeet [29–31], it is tempting to speculate that endfeet may be heterogeneous and display different local proteomes depending on their biochemical and cellular environment. In particular, it seems likely that endfeet in contact with capillaries perform different functions than those in contact with arterioles [7]. Successful molecular dissection of endfoot functional heterogeneity and plasticity is thus a promising venue for future research and will be instrumental in refining our understanding of the gliovascular unit and its diversity in health and disease.

In conclusion, our model proposes new interpretations of the role of astrocytes in NVC and suggests that the endfoot compartment, which enwraps blood vessels, could be a critical local mediator of NVC. Future research will be essential to characterize the inter-cellular and intra-cellular variability of these sub-cellular units in various physiological conditions and may provide cues for novel therapeutic strategies targeting NVC deficiencies.

## Acknowledgments

The authors thank the teams of Martine Cohen-Salmon and Blanca Diaz-Castro for sharing omics datasets and providing critical insights into local expression in astrocyte endfeet, as well as Prof. Keith Murai for kindly sharing unpublished endfeet FIB-SEM datasets. The authors have no competing interests to declare that are relevant to the content of this article.

## Data availability

The script that queries and aggregates the data from the four astrocyte omics studies used to model astrocyte-mediated NVC is available at https://gitlab.inria.fr/florian.dupeuble/endfeet omics analysis. The code of the model, implemented in Python, is available at https://gitlab.inria.fr/florian.dupeuble/astrovascular.

